# Thyroid hormone levels associate with insulin resistance in obese women with metabolic syndrome in Saudi Arabia: A cross-sectional study

**DOI:** 10.1101/595884

**Authors:** Manal Abdulaziz Binobead, Nawal Abdullah Al Badr, Wahidah Hazzaa Al-Qahtani, Sahar Abdulaziz AlSedairy, Tarfa Ibrahim Albrahim, Maha Hussain Alhussain, Tahani Ali Aljurbua, Shaista Arzoo, Wedad Saeed Al-Qahtani

## Abstract

**Background:** The obesity epidemic is a pressing global health concern, as obesity rates continue to climb worldwide. The current study was aimed mainly to evaluate the correlation between thyroid hormones and homeostatic model assessment of insulin resistance in Saudi obese women with metabolic syndrome.

**Methods:** 100 obese women aged 25 to 55 years were clinically evaluated, from which 72 women were diagnosed with the metabolic syndrome and 28 without metabolic syndrome. Insulin resistance was quantified using the homeostatic model assessment of insulin resistance method and the resulting values were analyzed for association with demographic, clinical, and metabolic parameters.

**Results:** This analysis revealed that body mass index, systolic blood pressure, and biochemical parameters and fasting insulin showed statistically higher levels in the group with metabolic syndrome compared to the group without metabolic syndrome. Similarly, values of waist circumference, fat ratio, cholesterol, free thyroxine, free triiodothyronine and homeostatic model assessment of insulin resistance results were higher in the group with metabolic syndrome as compared to the group without metabolic syndrome. Correlation analysis revealed positive association of thyroid-stimulating hormone with waist circumference (P=0.01), total cholesterol (P=0.002), fasting insulin (P=0.03) and homeostatic model assessment of insulin resistance results (P<0.01), and negatively associated with diastolic blood pressure (P=0.013) and age (P=0.05). Free thyroxine was positively associated with triglyceride level (P=0.003) and negatively associated with homeostatic model assessment of insulin resistance values (P=0.035) and fasting insulin. Free triiodothyronine was positively associated with body mass index (P=0.032) and waist circumference (P= 0.006) and negatively with age (P=0.004) and total cholesterol (P=0.001).

Homeostatic model assessment of insulin resistance test revealed elevated level with positive association of body mass index, waist circumference, biochemical parameters and thyroid-stimulating hormone in insulin resistant obese women. Higher level of free triiodothyronine was found to be associated with low insulin sensitivity.

## Introduction

The obesity and overweight are associated to the risk of health complications and premature death, obesity is the greatest contributing factor underlying the metabolic syndrome (MetS) (Danaei et al., 2014). MetS is a chronic medical condition manifested by a cluster of symptoms (e.g., low high-density lipoprotein cholesterol (HDL-C) levels, high blood pressure (BP), high triglyceride (TG) levels, insulin resistance (IR), and other anthropometric and biochemical factors) that are associated with developing cardiovascular disease and type 2 diabetes mellitus (Ford et al., 2005). A precursor to type 2 diabetes. IR basically refers to the inability of insulin to perform its function at the optimum concentration required for its biological activity (Harris et al., 1998; Ferrannini, 2004). This causes responsible for this inability can range from defective glucose output in the liver to impaired insulin uptake in the muscle (Farasat et al., 2011).

Healthy thyroid activity is required to maintain the overall health of an individual. Several studies have described the effect of thyroid hormones on body mass index (BMI). Hypothyroidism leads to weight gain, while hyperthyroidism causes weight loss (Hoogwerf and Nutall, 1984). Moreover, it has also been established that obesity affects thyroid gland function (Topsakal et al., 2012). Previous studies have associated thyroid hormones with insulin activity, toward regulating the metabolism of glucose; the dysregulation of this pathway contributes to IR (Ravi and Gokaldas, 2015). Thyroid hormones regulated a variety of proteins involved in maintaining insulin sensitivity (Klieverik et al., 2009). The loss of insulin sensitivity, IR, is associated with obesity and has been used as a predictor of developing cardiovascular disease and type 2 diabetes (Naslund et al., 2000). Multiple studies have evidenced the cooperative relationship between thyroid hormones and insulin in glucose metabolism (Lacobellis et al., 2005; Chakarabarti et al., 2007). Maintaining glucose homeostasis involves the complex interplay between physiological pathways that regulate insulin secretion and modulate its activity (Farasat et al., 2011). The American Association of Clinical Endocrinologists (AACE) has provided guidelines for the diagnosis of abnormal thyroid function and for the treatment of thyroid dysfunction in patients with abnormal serum levels of thyroid-stimulating hormone (TSH) (Gharib et al., 2004). The possible role of TSH in adipogenesis and IR has already been established (Bastemir et al., 2007). Among its various metabolic effectors, WC and BMI correlate positively with serum TSH levels (Knudsen et al., 2005); however, the relationship between IR and TSH remains largely unexplored, particularly in the Saudi, female population affected by MetS.

Therefore, the aim of this cross-sectional study is to identify associations between TSH, IR, and other clinically-relevant metrics in obese women with and without MetS in Saudi Arabia. The objectives of this study include: 1) to evaluate the anthropometric, clinical, and biochemical characteristics of the study subjects; 2) to analyze these characteristics for correlations with insulin sensitivity level; 3) to detect associations between TSH levels and the subjects’ clinical and biochemical characteristics, MetS diagnosis, and insulin sensitivity level.

## Materials and Methods

### Study subjects

The analysis was carried out on 163 obese and overweight women aged 25 to 55 years. All of the patients had BMI ≥ 25 kg/m^2^. The presence of medical conditions was assessed through self-report. A pre-structured and pre-tested questionnaire were used to gather demographic information and personal and family medical history. Informed consent was obtained from all participants.

#### Inclusion criteria

This study included women aged 25 to 55 years with BMIs over 25 kg/m^2^ (Fig. 1).

**Figure 1.**
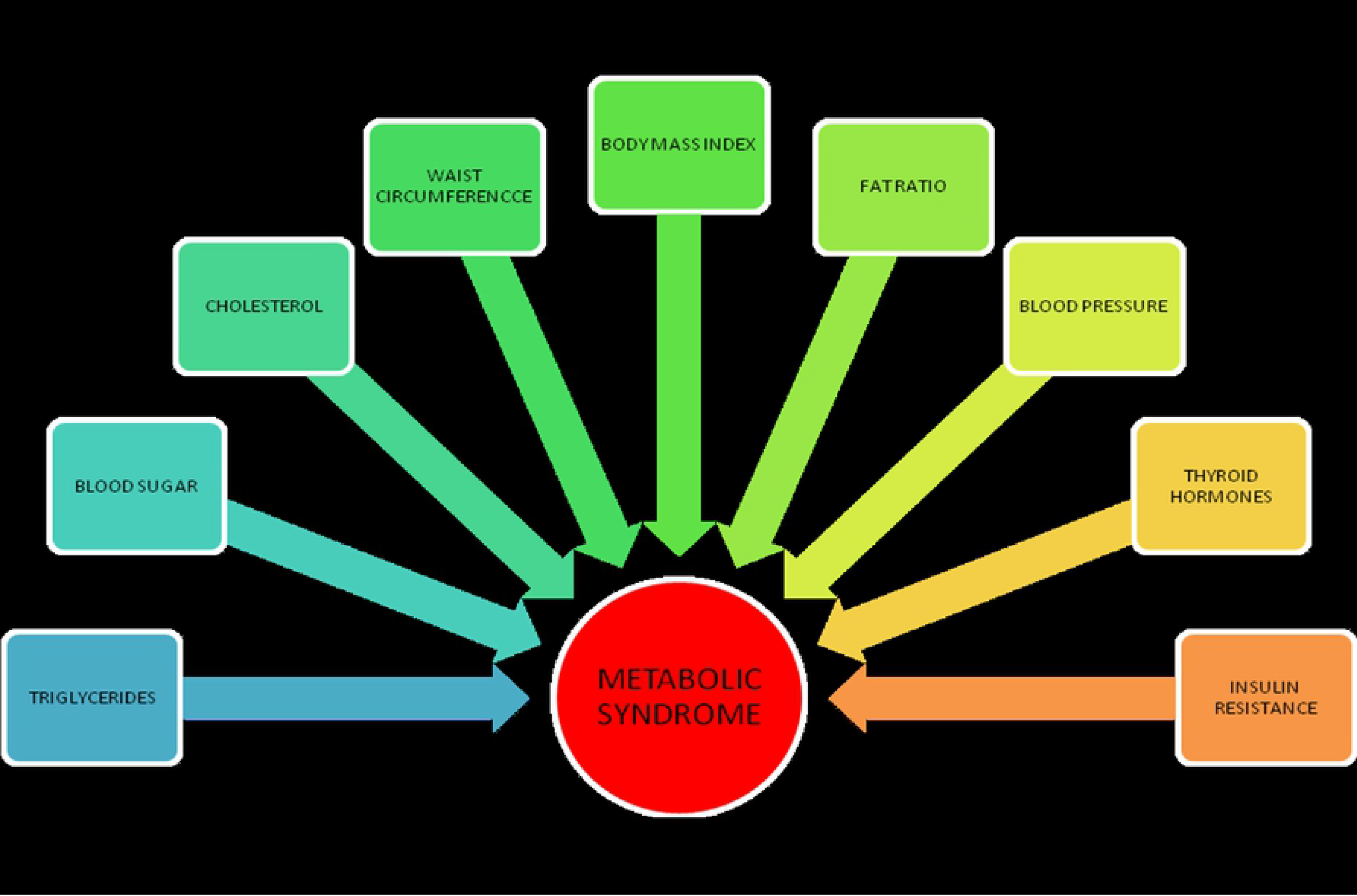
Flowchart schematic of study subject selection using the inclusion and exclusion criteria

#### Exclusion criteria

Subjects with a history of smoking, polycystic ovary syndrome, chronic renal failure, thyroid disease, chronic hepatopathy, or cancer as well as subjects taking antihypertensive drugs and statins, contraceptive drugs, hormone replacement therapy, any medications known to interfere with glucose and/or insulin secretion and/or metabolism were excluded from the study (Fig. 1).

#### Demographic data

A pre-structured and pre-tested questionnaire was used to gather self-reported demographic information and individual and familial medical history.

#### Ethical considerations

Informed consent was orally obtained from all participants before they gave voluntary consent for this study and approved by IRB.

### Anthropometric measurements

Anthropometric measurements were carried out three times by a single tester.

#### Height and weight

Height was measured without shoes and using a stadiometer. Body weight was morning determined in lightweight clothing, with a digital scale.

#### Body mass index (BMI)

BMI was calculated as the weight (kilograms) divided by the square of height (meters).

#### Waist circumference (WC)

Subject’s WC was measured using a flexible measuring tape, midway between the xiphoid and the umbilicus during the mid-inspiratory phase.

#### Blood pressure (BP)

Two BP measurements were taken with the subject in the seated position at a 2- to 3-minute interval, after resting for at least 15 minutes. The average of these two readings was used for all patients.

### Biochemical parameters

Blood samples were drawn after an overnight fast. Serum samples were analyzed for fasting blood glucose (FBG), TG, total cholesterol (TC), low-density lipoprotein cholesterol (LDL-C) and high-density lipoprotein cholesterol (HDL-C) using commercially-available kits (Beckman-Coulter, CITY, STATE, USA). Serum insulin concentration was determined using an electrochemiluminescence-based assay (Immulite 2000, CITY, STATE, USA). Serum FT_4_, FT_3_, and TSH levels were also determined by electrochemiluminescence-based immunoassay (Roche Diagnostics, CITY, Germany).

The homeostatic model assessment (HOMA) ratio formula was used to quantify IR (Mathews et al., 1985).

HOMA-IR = *[fasting plasma insulin (μIU/ml) × fasting plasma glucose (mmol/l)] / 22.5*

A HOMA-IR cut-off value chosen was 2.7 (> 2.7 resistant, < 2.7 sensitive).

### Diagnosis of metabolic syndrome (MetS)

MetS was diagnosed according to standard protocol (Grundy et al., 2005) based on the presence of the following criteria: 1) TG ≥ 150 mg/dL; 2) LDL-C < 130 mg/dL; 3) HDL-C < 40mg/dL; 4) TC < 200 mg/dL; 5) FBS ≥ 100 mg/dL; 6) SBP ≥ 130 mmHg; 7) DBP ≥ 85 mmHg; 8) WC > 80 cm; 9) TSH > 2.5 IU/mL. Subjects with levels over the cut-off values were considered as MetS+ and subjects with levels under the cut-off values were considered to be MetS−.

### Statistical analysis

All data were analyzed using the Statistical Package for the Social Sciences (SPSS) software package v25 (IBM, Chicago, IL, USA). Statistical comparisons between the MetS+ and MetS− groups were achieved with the one-way analysis of variance (ANOVA). Significance assessments were carried out using Duncan’s new multiple range test. Values are expressed as means and standard deviations. We used hierarchical cluster analysis to assess the relationship between TSH levels and HOMA-IR values. Each experiment was repeated at least three times. A *P*-value of less than 0.05 was regarded as statistically significant.

## Results

Of the 163 individuals considered, 63 were eliminated based on exclusion criteria (Fig. 1). The study population consisted of the remaining 100 obese (BMI > 25 kg/m^2^) women aged 25 to 55 years. Of these, 72 women were diagnosed as MetS+ and the remaining 28 were MetS−. The anthropometric and biochemical characteristics are presented in Table 1. BMI, SBP, TC, TG, HDL-C, FBP, TSH, and fasting insulin levels were statistically higher in the MetS+ group than the MetS− group. Similarly, the values for WC, fat ratio, LDL-C, FT_4_, FT_3_, and HOMA-IR were higher in the MetS+ group than the MetS− group; however, these differences were not statistically significant.

**Table 1.**
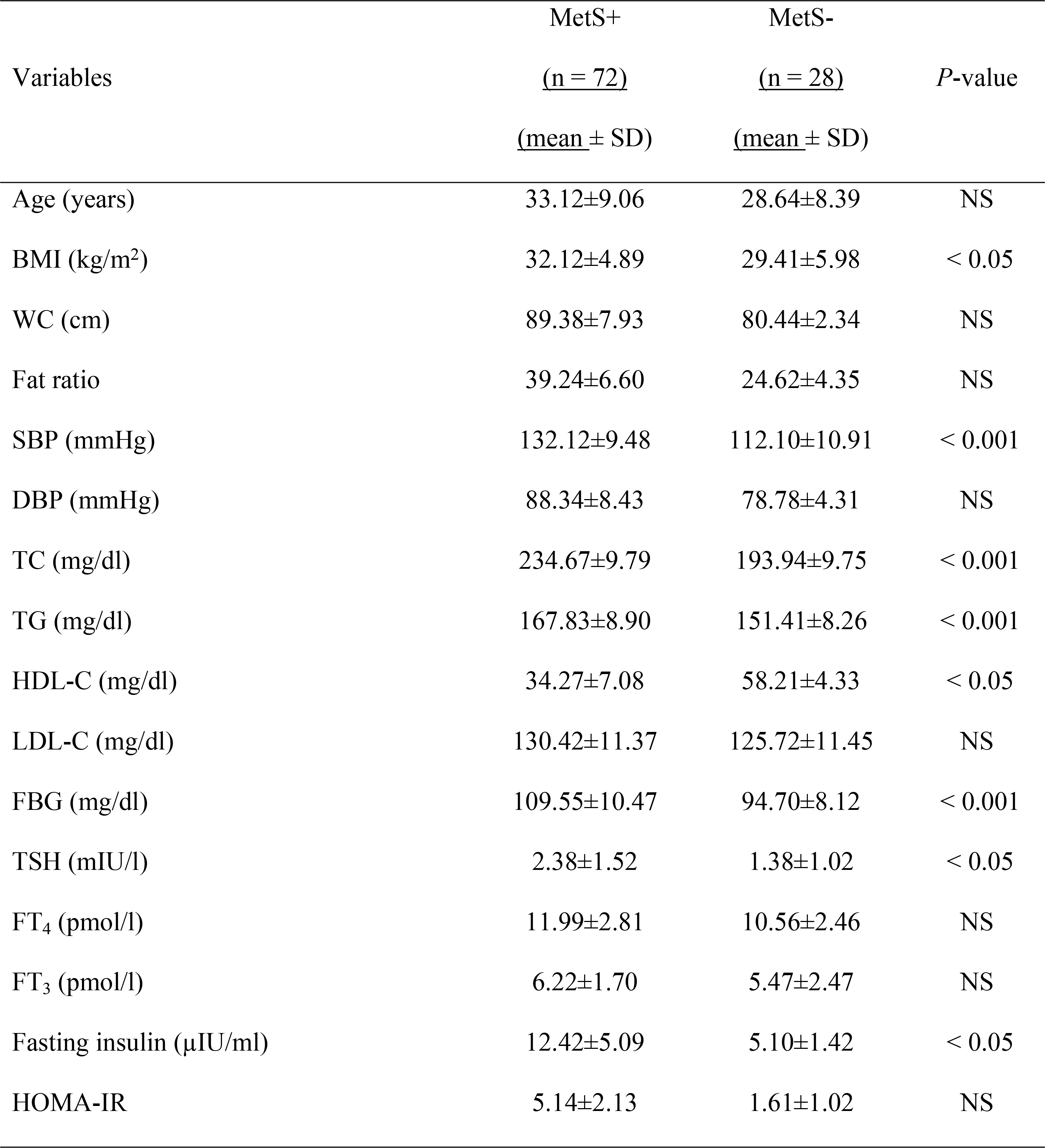
Anthropometric and biochemical characteristics (data are given as a mean and standard deviation)

Based on the Pearson’s correlation coefficients (Table 2), TSH positively associated with WC (*P* = 0.01) and TC (*P* = 0.002) and negatively associated with diastolic blood pressure (DBP) (*P* = 0.013) and age (*P* = 0.05). TSH also positively associated with insulin (*P* = 0.03) and HOMA-IR (*P* < 0.01). FT_4_ positively associated with TG level (*P* = 0.003) and with HOMA-IR value (*P* = 0.035). FT_3_ positively associated with BMI (*P* = 0.032) and WC (*P* = 0.006) and negatively with age (*P* = 0.004) and TC (*P* = 0.001).

The comparison between insulin-sensitive and -resistant women in terms of the clinical and metabolic characteristics is presented in Figure 2. Using a cut-off value of 2.7 for HOMA-IR (> 2.7 resistant, < 2.7 sensitive), BMI, WC, TC, TG, LDL-C, FBS, and TSH were higher in the resistant group than the sensitive one. The positive association between IR and BMI (*P* < 0.001) and WC (*P* < 0.05) was statistically significant. Similarly, TSH was significantly associated with IR (*P* = 0.03). Higher FT_3_ level associated with low levels of insulin sensitivity.

**Figure 2.**
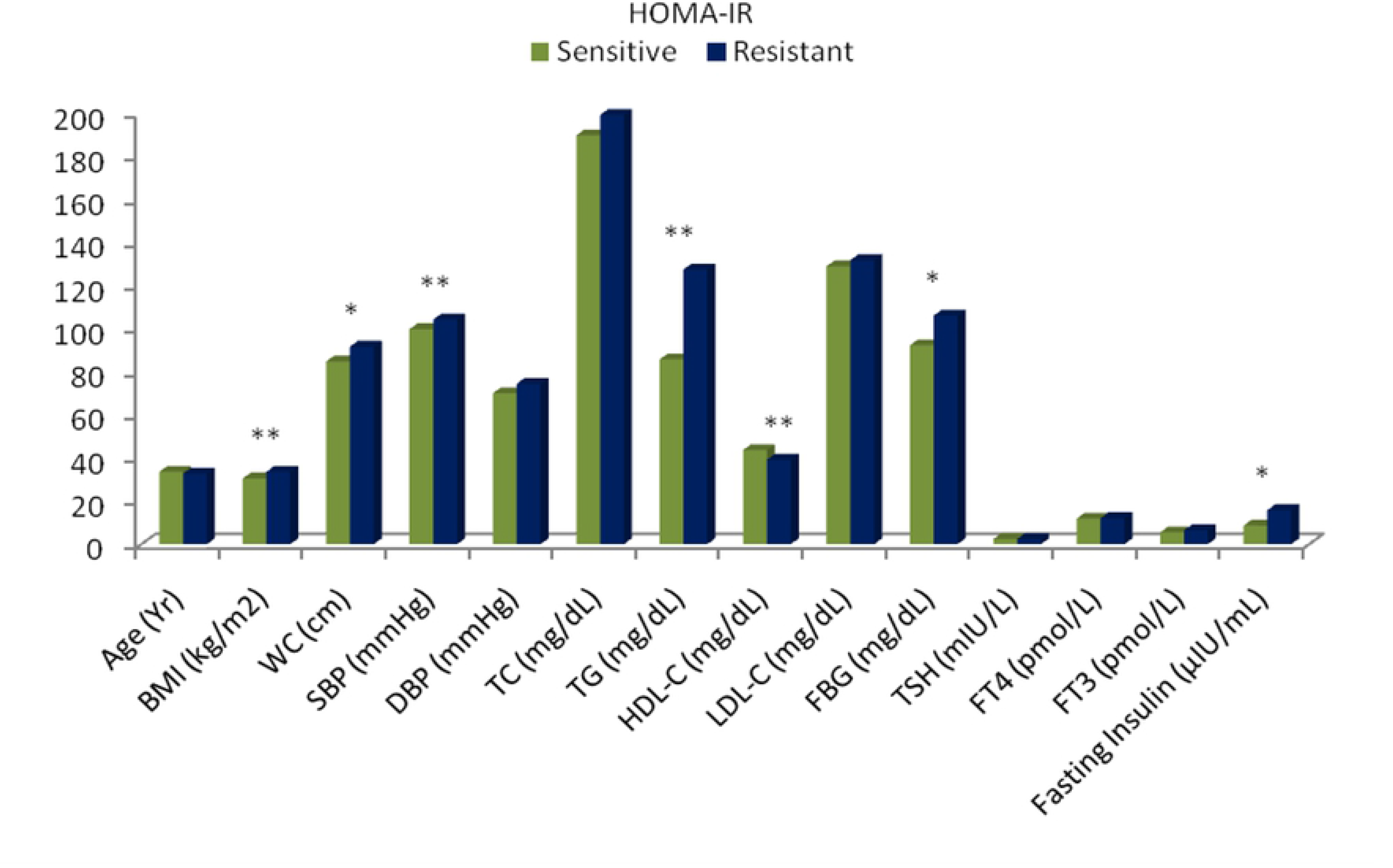
Comparison of clinical and metabolic characteristics according to HOMA-IR. Insulin-sensitive (green, n = 21) and insulin-resistant (blue, n = 51) obese women. \**P* < 0.05, \*\**P* < 0.001. BMI = body mass index, WC = waist circumference, SBP = systolic blood pressure, DBP = diastolic blood pressure, TC = total cholesterol, HDL-C = high-density lipoprotein cholesterol, LDL-C = low-density lipoprotein cholesterol, FBG = fasting blood glucose, TSH = thyroid-stimulating hormone, FT_4_ = free thyroxine, FT_3_ = free triiodothyronine.

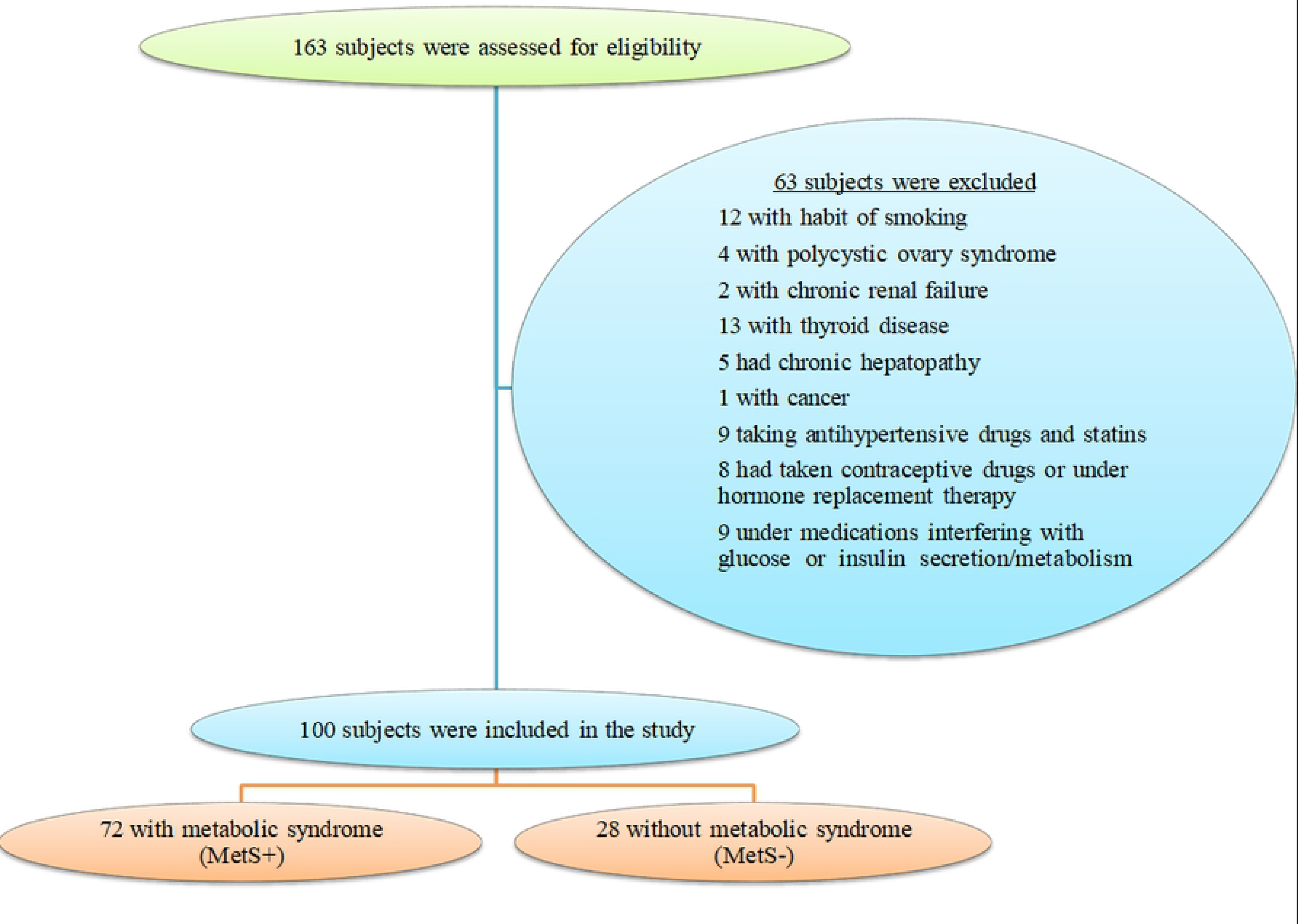

Furthermore, hierarchical cluster analysis grouped the clinical and metabolic data according to HOMA-IR values and revealed statistically significant associations between these groups (Fig. 3).

**Figure 3.**
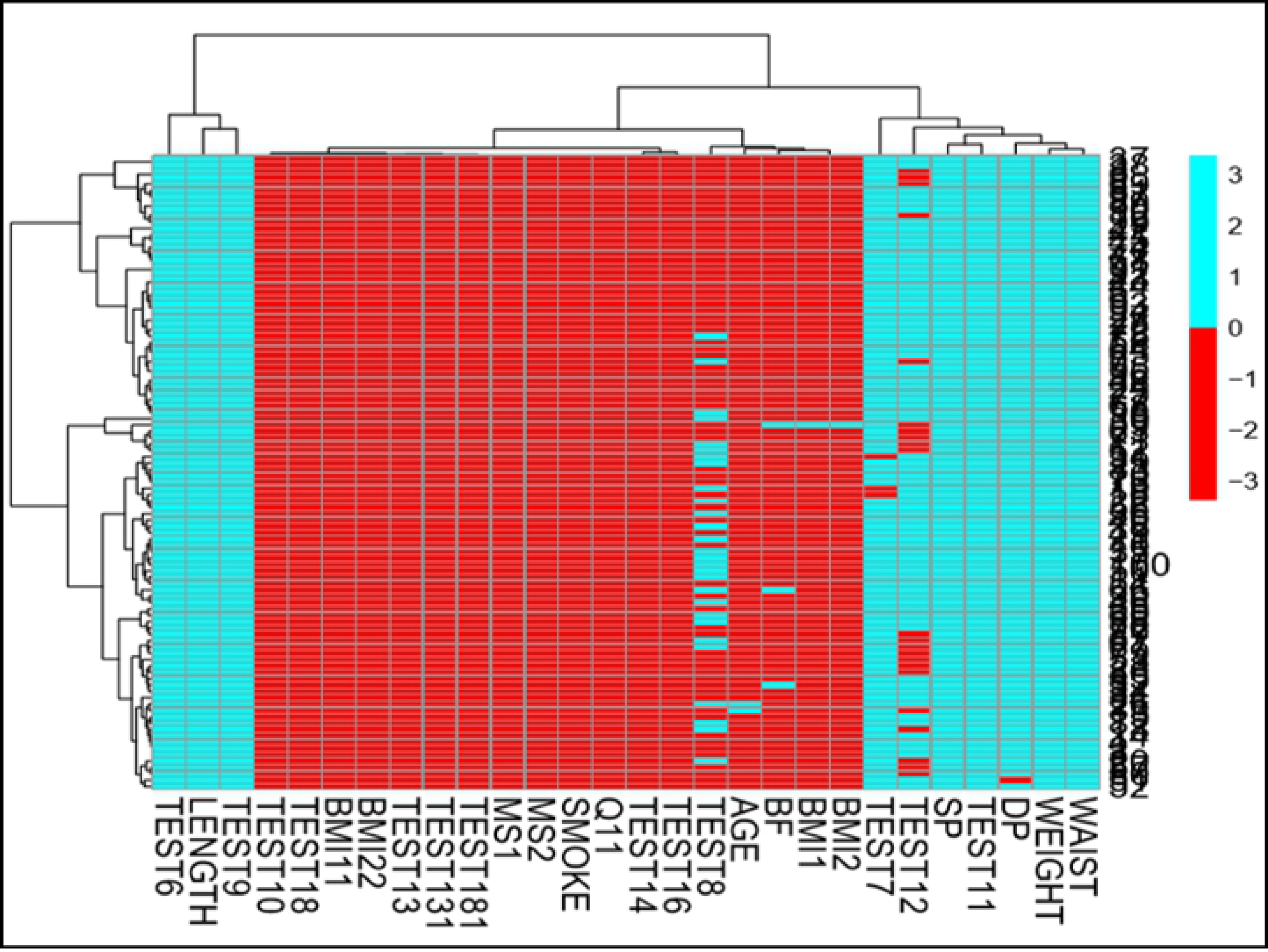
Hierarchical cluster analysis showing a significant correlation between HOMA-IR values and clinical and metabolic characteristics between both groups.

**Table 2.**
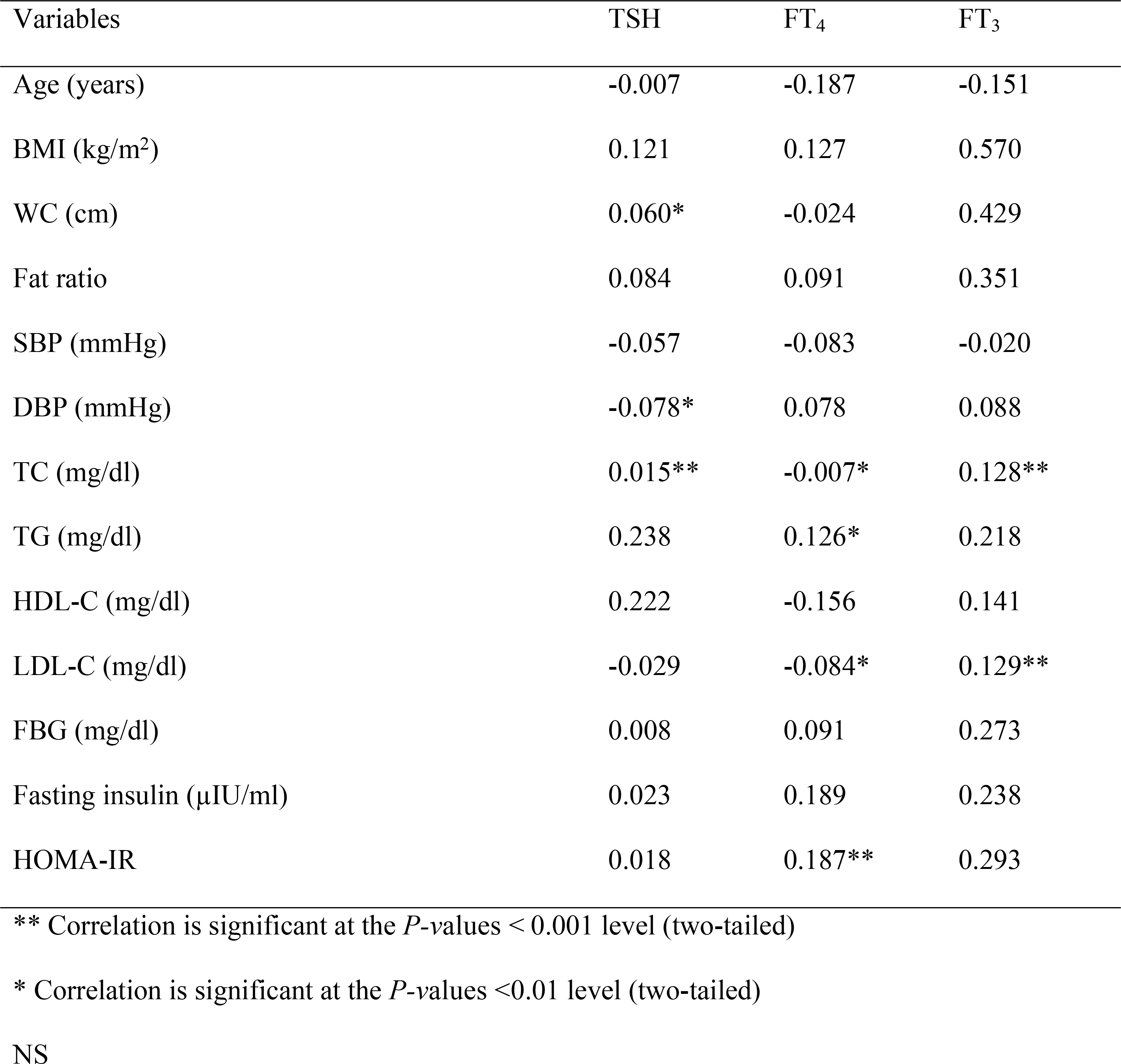
Pearson correlation coefficients (r) of TSH, FT_4_, and FT_3_ with demographic, anthropometric, and MetS-associated factors

## Discussion

MetS describes a constellation of different metabolic irregularities, which are often associated with thyroid hormones and IR. The present study investigated the relationships between thyroid hormones and IR in obese women diagnosed with MetS in Saudi Arabia. It is common knowledge that excessive weight gain and the resulting obesity increases the likelihood of incurring MetS. One of the risk factors for IR and hyperlipidemia is hypothyroidism. Hypothyroidism is associated with weight gain and concomitant changes to the other components that comprise MetS (Tarcin et al., 2012).

The present analysis established that HOMA-IR and TG values are comparatively higher in women that are MetS+ relative to their MetS− counterparts. Moreover, TSH was found to positively associate with WC and total cholesterol levels. The finding is in line with previous studies (Pergola et al., 2007, Ayturk et al., 2009). Increases in TSH concentration, weight, and TC level are likely indicative of subclinical hypothyroidism. This result serves as evidence for the association between elevated TSH levels and obesity and MetS. Correlations between hypothyroidism and IR have been thoroughly established by several earlier studies (Singh et al., 2010; Pergola et al., 2007; Ravi and Gokaldas, 2015). The analysis described here revealed positive associations between FT_4_ and HOMA-IR values and fasting insulin levels. Low serum levels of free T_4_ were observed in MetS+ women. This strongly suggests that low FT_4_ levels mediate the development of IR. Thus, the association between thyroid hormone and HOMA-IR cannot be discounted and requires further investigation. In the present study, FT_4_ and TG levels correlated positively, which is in contrast to one described a previous study (Kim et al., 2009). Furthermore, a positive correlation was observed between FT_4_ and MetS-associated variables (Tarcin et al., 2012). Our results contradict the findings of Ayturk et al. (2009), who did not detect any correlation between free thyroid hormones and MetS. The increases in TC levels and LDL-C specifically are indicative of insulin sensitivity.

In the present study, increased FT_3_ levels positively associated with increases in BMI. This is in accordance with a previous finding where associations between free or total thyroid hormone levels and body weight and BMI increases were observed (Roef et al., 2012). Interestingly, the increase in FT_3_ concentration was independent of other metabolic parameters and insulin sensitivity; these results corroborate the findings of a previous study (Pergola et al., 2007). FT_3_, alone or in combination with insulin, regulates the uptake and breakdown of glucose levels. A positive correlation between TG levels and HOMA-IR values was observed in MetS+ subgroup in the present study. Some of the contradictory findings in the present study may stem from the study design or variations in the health status of the study subjects.

The HOMA approach is a reliable, time-tested method for quantifying IR that is both well established and regarded in the field. The HOMA-based analysis of MetS+ and MetS− obese women described here provides empirical evidence that BMI positively correlates with IR, which supports the findings of a previous, independent study (Geloneze et al., 2009). Moreover, the positive association between HOMA-IR values and elevated TG and total cholesterol levels nicely reflects its preponderance for MetS. Furthermore, the positive correlation between HOMA-IR values and TSH levels observed in the present study highlights the role played by the thyrotropin hormone in adipogenesis. These results are consistent with the findings of Bastemin et al. (2007) and were further validated using hierarchical clustering.

## Acknowledgements

The authors extend their appreciation to the Deanship of Scientific Research at King Saud University for funding this work through the Research Project No R6-17-03-65.

